# Context- and scale-dependent effects of thymol bioactivity on biological networks: contributions from quail under heat stress

**DOI:** 10.1101/2022.06.10.495659

**Authors:** Maria Emilia Fernandez, Rocío Inés Bonansea, Agustin Lucini Mas, María Verónica Baroni, Raul Hector Marin, Maria Carla Labaque, Jackelyn M. Kembro

## Abstract

A defining feature of healthy function is adaptability, the capacity to dynamically respond to novel and/or unpredictable factors, such as exposition to environmental challenges. Here we explore the role of context (i.e., order in which factors are applied) on the configuration of networks of behavioral and physiological traits from Japanese quail that were supplemented dietary thymol prior, jointly or after onset of a chronic heat stress protocol (i.e., supplementation strategies). Basal diet and standard environmental temperature were used as controls. We begin by showing that each supplementation strategy evolved differently over the 5-week experimental period in regard to body weight and feed-intake. At the end of this experimental period, context-dependency was also observed in the non-trivial functional relationships among 27 traits from 4 subsystems from distinct spatial-temporal scales. When considered separately, whole-organism level subsystems (i.e. somatic maintenance and egg production traits), are predominantly functionally related to environmental temperature. Conversely, at the molecular level, the liveŕs antioxidant system response is fundamentally dominated by the supplementation strategy. Interestingly, the serum’s antioxidant system shows an intermediate response, not dominated by a given factor. Overall, network configurations were highly dependent on context, and could be associated with specific induced physiological states. Our work constitutes the first study that includes network, integrative, and experimentally comparative analyses applied to the field of dietary supplementation under environmental challenges. This perspective could help understand complex biological responses important for developing efficient and welfare orientated supplementation protocols for farm animals and for interpreting the background in this field.

## Introduction

A defining feature of healthy function is adaptability, the capacity to continuously respond to novel and/or unpredictable stimuli [1] such as exposition to environmental challenges. Specifically, biological response patterns to external stimuli can involve changes in a large number of traits observable over a broad range of spatio-temporal scales, ranging from the whole-organism to the molecular scale [2, 3]. Since these traits are functionally related to each other, the nonlinear relations among them [4, 5] can be portrayed as a network (or network of networks) as shown in Figure 1. Noteworthy, the topology and link strength between the nodes of the resulting physiological network is dynamic, and can change depending on the physiological state (e.g. health vs. disease; light vs. deep sleep [6, 7]) of the animal. Moreover, these networks require energy input through metabolism and coordination of many processes in order to fulfill the energy needs of all cells [8]. Since resources for maintaining physiological integrity are limited, diverging resources to respond to an environmental challenge can result in potentially compensating impacts on other subsystems [9–12]. Hence, prolonged environmental challenge may result in observable dynamical effects on the underlying physiological network topology and system connectivity.

**Fig 1.**
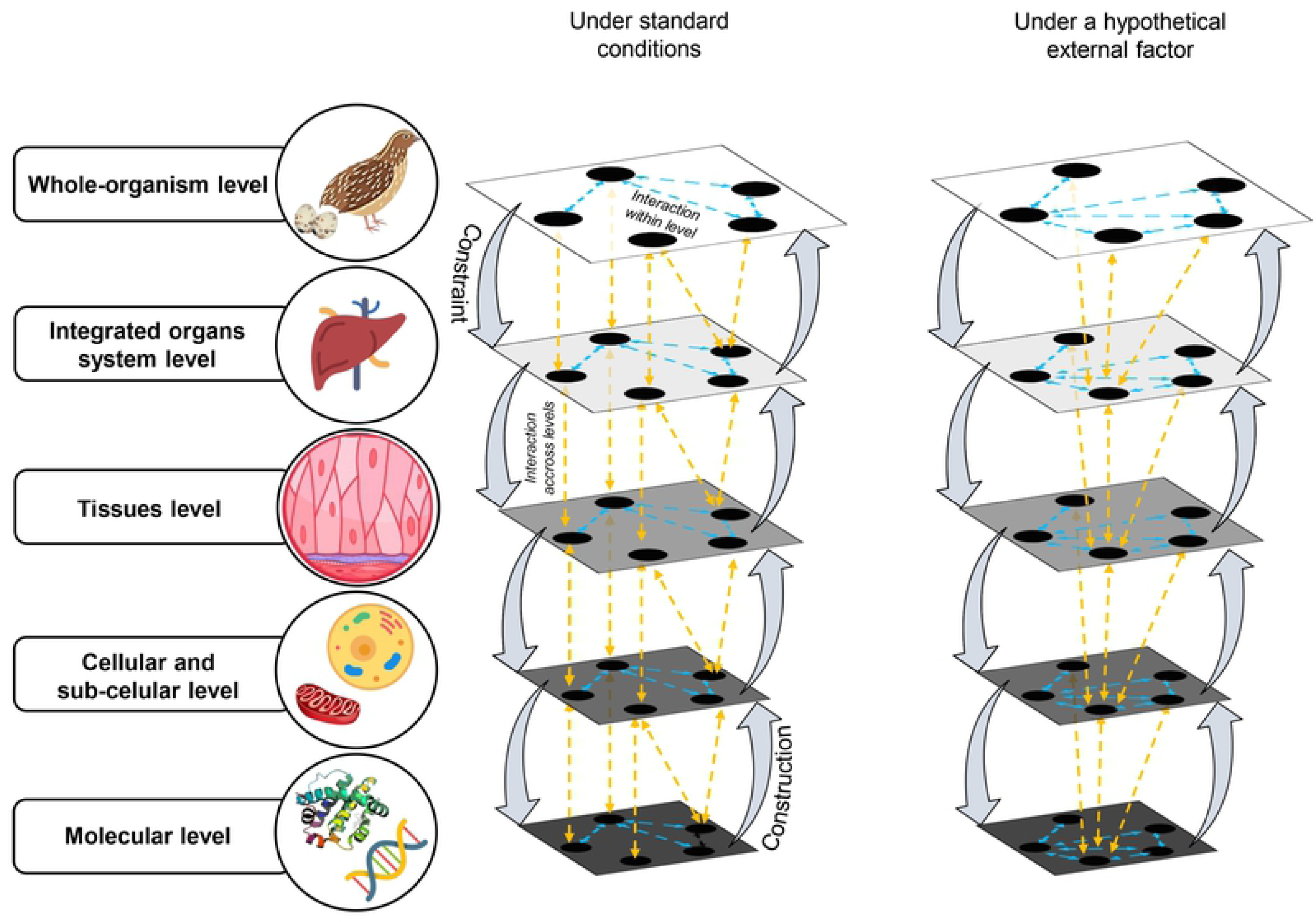
Network perspective for understanding living organisms’ response to stimuli and emergence of physiological states (this figure has been adapted from figures 1 and 4 of references [13] and [4], respectively).

Black circles (i.e., nodes of the network) in each layer represent components inherent to a specific spatio-temporal scale (level). Bidirectional arrows with dotted line (i.e., the edges of the network) within and between layers indicate the simultaneity of action (causation) among components and levels. Downward and upward lateral tick arrows also represent causation. Downward causation is the set of constraints imposed by the higher levels of the dynamics at lower levels through determining many of the initial and boundary conditions of the system [4]. Upward causation is the set of processes by which the lower elements in a system interact and produce changes at higher levels [4]. Thus, physiological levels interact both horizontally (within the level) and vertically (across levels) through nonlinear rules (i.e., circular causality), and distinctive physiological states emerge from specific configurations of this network of networks. Figure was designed using images from freepik.com (Freepik Company S.L.), flaticon.es (Freepik Company S.L.), shutterstock.com (Shutterstock, Inc.). Attribution to images’ authors under the license terms is detailed after the references section.

Although complex nonlinear responses, networks and resource allocation are all fundamental general premises in biology, they most often are not specifically taken into account. This is particularly noticeable in studies evaluating the effects of environmental challenges and counteracting measures in farm animals. For example, regarding dietary supplementation strategies, recent reviews [14–17] have shown that most research in this field has focused on “improving” traits that are considered important, thus, predominantly relaying on traditional univariate analysis (i.e., statistical analysis of each variable separately). However, considering the above-described multivariate nonlinear dynamical nature of networks (Fig 1), a more comprehensive approach that includes the analysis of the functional relationships among traits is necessary in order to understand the underlying mechanisms behind complex and often contradictory response patterns observed in this field [14–17].

As with other environmental challenges, elevated environmental temperature in poultry species [14,18–20] has broad effects on different subsystems, spanning from impairing resource allocation for somatic maintenance (e.g. feed intake, weight gain) and egg production [21], modulation of hypothalamic centers and neuroendocrine axis to cellular energetics and genetic regulation [22, 23], and, in the long-term, antioxidants depletion [14,23–27]. Given the impact of heat stress in the productivity of poultry as well as other farm species [18,19,28], phytochemical compounds, such as thymol (2-isopropyl-5-methylphenol), have been proposed as a strategy for counteracting its consequences. This contention has been mainly based on their antioxidant properties [14] as well as other potentially beneficial effects *in vivo* [14,29–32]. Specifically, thymol can act directly, as a free radical scavenger [33–35], and indirectly, upregulating Nrf2 conducive to increased synthesis of antioxidant enzymes [36, 37]. In mammals and birds [16,38–41] it also can increase polyunsaturated fatty acids concentration and decrease lipid peroxidation in several tissues and in the egg yolk, and it can promote beneficial whole-organism effects associated with somatic maintenance (e.g. body weight, behavior, feed intake) and egg production performance [16,42–44].

As can be noted from the above descriptions, through various mechanisms and biochemical pathways, both thymol supplementation and heat stress have the capacity to influence some of the same traits (e.g., antioxidant system, somatic maintenance). Moreover, both factors separately and together have time-dependent effects on traits such as feed intake [21, 45], and antioxidant enzymes synthesis. For example, short-term exposure to thymol in fish [36] can induce a pro-oxidant effect with a decrease in antioxidant enzymes synthesis, while, in the long-term the contrary has been observed in mice [37]. In poultry [46–49] short-term exposition to heat stress induces the synthesis of antioxidant enzymes though Nrf2, while in the long term promotes antioxidants depletion including enzymes [50, 51]. Therefore, as a contingency of the transition between short- and long-term response to each of these two factors, the outcome of combining them is non-trivial and could be reliant on the order in which exposure to them occurs. Moreover, the dynamical responses to each factor may overlap or alter the trajectory of the response to the other factor. Hence, context (i.e., order in which factors are applied) is fundamental in understanding both the dynamical response to chronic heat stress and dietary supplementation, as well as the underlying physiological state that emerges after prolonged exposition.

Precedent studies in poultry and other farm animals have mainly focused on experimental designs aimed at evaluating one of two types of supplementation strategies. Specifically, those in which thymol or other phytogenic supplementation was initiated either prior to (appealing for potential preventive effects) or simultaneously with heat stress exposition [14,45,52–54]. For example, adult Japanese quail exposed to chronic cyclic heat stress and previously supplemented with thymol, showed a heterophil/lymphocyte ratio similar to their control non-stressed counterparts [53]. Similarly, in growing broilers simultaneously subjected to heat stress and thymol supplementation, beneficial effects on body weight gain, feed conversion ratio, water intake and serum glutathione peroxidase were found in comparison to non-supplemented controls [54]. Interestingly, experiments in birds and rodents under other environmental challenges that induce oxidative stress have also assessed a third type of supplementation strategy in which thymol administration began after exposure to the challenge [55–58], appealing to a potential palliative effect. However, to our knowledge, these three supplementation strategies have not been compared yet in a single experiment in any species.

Herein, we hypothesize that the potential for thymol to modulate heat stress impact on key traits from different subsystems and spatio-temporal scales is dependent on the moment (time-schedule) in which supplementation initiates with regard to the beginning of heat stress exposure. We begin by studying the evolution of the system over the 37-day experimental period through monitoring body weight and feed intake in female Japanese quail that were supplemented with dietary thymol prior, jointly or after the onset of a chronic heat stress protocol (i.e., supplementation strategies). Then, we assessed whether the differential trajectories observed under each supplementation strategy translated into specific network configurations. To this end, a combination of analytical techniques (i.e., clustering, Pearson correlations and network analysis) were applied to 27 key traits associated with 4 subsystems representing scales ranging from whole-organism to molecular. Our analytical approach, aimed at understanding the relationships among key traits under different contexts, provides a first approximation of how the underlying physiological state of the animal can be modulated by environmental challenges within the framework of complex nonlinear responses, networks, and resource allocation. Moreover, this approach could be also expanded to other multivariate studies in the field of dietary supplementation and stress, further enhancing knowledge on emergent complex responses in farm animals.

## Materials and methods

### Ethical statement

All experimental procedures were in compliance with the *Guide for the Care and Use of Laboratory Animals* issued by the National Institutes of Health [59]. The experimental protocol was approved by the Institutional Committee for the Care and Use of Laboratory Animals (Comité Institucional para el Cuidado y Uso de Animales de Laboratorio (CICUAL) of the Facultad de Ciencias Exactas, Físicas y Naturales, Universidad Nacional de Córdoba (ACTA 4/2015 Resolución 571-HCD-2014) and the Instituto de Investigaciones Biológicas y Tecnológicas-Consejo Nacional de Investigaciones Científicas y Técnicas (Acta 27, IIByT’s Board of directors). The raw data of all variables presented herein is publicly available on Figshare (private link: https://figshare.com/s/d13f182242487f522376).

### Animals and husbandry

Husbandry of female Japanese quail (*Coturnix japonica*) was performed according to laboratory routines described elsewhere [41, 43]. Briefly, at 28 days of age, females were housed in pairs in enriched cages measuring 20cm × 45cm × 25cm (length × width × height), with 50% of the floor covered by corrugated cardboard which was renewed weekly [60–62]. Two feeders and an automatic nipple drinker were positioned in each cage. At all stages feed and water were provided *ad libitum*. The photoperiod was 14 Light:10 Dark (being the light period from 6 h to 20 h; approximately 300-320 lx). The environmental temperature was maintained at 24 ± 2°C.

### General procedure

Sixty-four, 96-days of age, female quail with body weights within the range of 161-213 g and in their egg-laying peak (5-7 eggs/week) were obtained from a larger population. These females were randomly reallocated in pairs and distributed in different experimental rooms depending on the environmental temperature they would be exposed to, as described in previous protocols of our laboratory [53]. As shown in Figure 2, all birds continued receiving a basal diet and were maintained at standard environmental (room) temperatures until testing began. The first day of the experimental period, females were randomly assigned to a control basal diet (Basal; 16 birds) or to a chronic dietary supplementation with 6.25 g of thymol/kg of feed (see preparation in S1 Appendix) that was supplied accordingly to one of three strategies Prior, Joint or Post strategy (16 birds in each strategy, see details described in the following paragraphs). At day 14 of the experimental period, half of the birds within each of these dietary groups were maintained at standard environmental temperatures (no heat stress; NHS, 23.6 ± 0.1°C) or were submitted to a chronic cyclic heat stress (HS) protocol induced by daily increasing the environmental (room) temperature from 23.6 ± 0.1°C to 34.2 ± 0.1°C between 8-17 h, over a 23-day period, according to [45] (see details in S2 Appendix). Finally, at day 37 of the experimental period, birds were euthanized. In order to minimize the number of birds euthanized, data from the same Basal-NHS or Basal-HS birds were used for statistical comparisons within each supplementation strategy. Each supplementation strategy was defined as follows:

a. Prior strategy: thymol supplementation started the first day (day 1) of the experimental period and continued for 5-weeks (37 days), up to the end of the experimental period (Fig 2). Birds under these conditions will be referred to as “Prior-NHS” or “Prior-HS” depending whether they were subjected to standard environmental temperature or HS, respectively. Given that supplementation begins prior to the onset of HS, in these birds the supplementation period encompassed the first 14 days under standard temperature plus the following 23 days under HS. We have previously shown that thymol supplementation in a dose identical to that used herein, when administered for a threshold period of 14 days, promotes favorable changes in the fatty acids profile of quail egg yolk [41, 63] and is also conducive to a plateau of thymol concentration in yolk and droppings, with no negative effects on female performance [63].
b. Joint strategy: thymol supplementation started the same day as the HS protocol, on the fourteenth day (day 14) of the experimental period, and continued for 23 days up to the end of the experimental period (Fig 2). Birds under these conditions will be referred to as “Joint-NHS” or “Joint- HS”, depending on the environmental temperature they were exposed to. This supplementation strategy with thymol has previously shown beneficial influence on physiological and behavioral variables of adult female quail [45]. Additionally, as stated in the introduction, supplementation protocols in a joint strategy-like fashion are the most frequently found in literature regarding supplementation with phytogenic compounds in poultry under heat stress [14,52–54].
c. Post strategy: thymol supplementation started the twenty-third day (day 23) of the experimental period and continued for 14 days, up to the end of the experimental period (Fig 2). These birds will be referred to as “Post-NHS” or “Post-HS” depending on the environmental temperature they were exposed to. Note that in Post-HS the onset of supplementation corresponds to 9 days after HS was began (Fig 2). In this manner, supplementation period encompassed the last 14 days of HS. This 9-day heat stress protocol has been previously used in immune-neuro-endocrine modulation studies in quail [23, 53].

**Fig 2.**
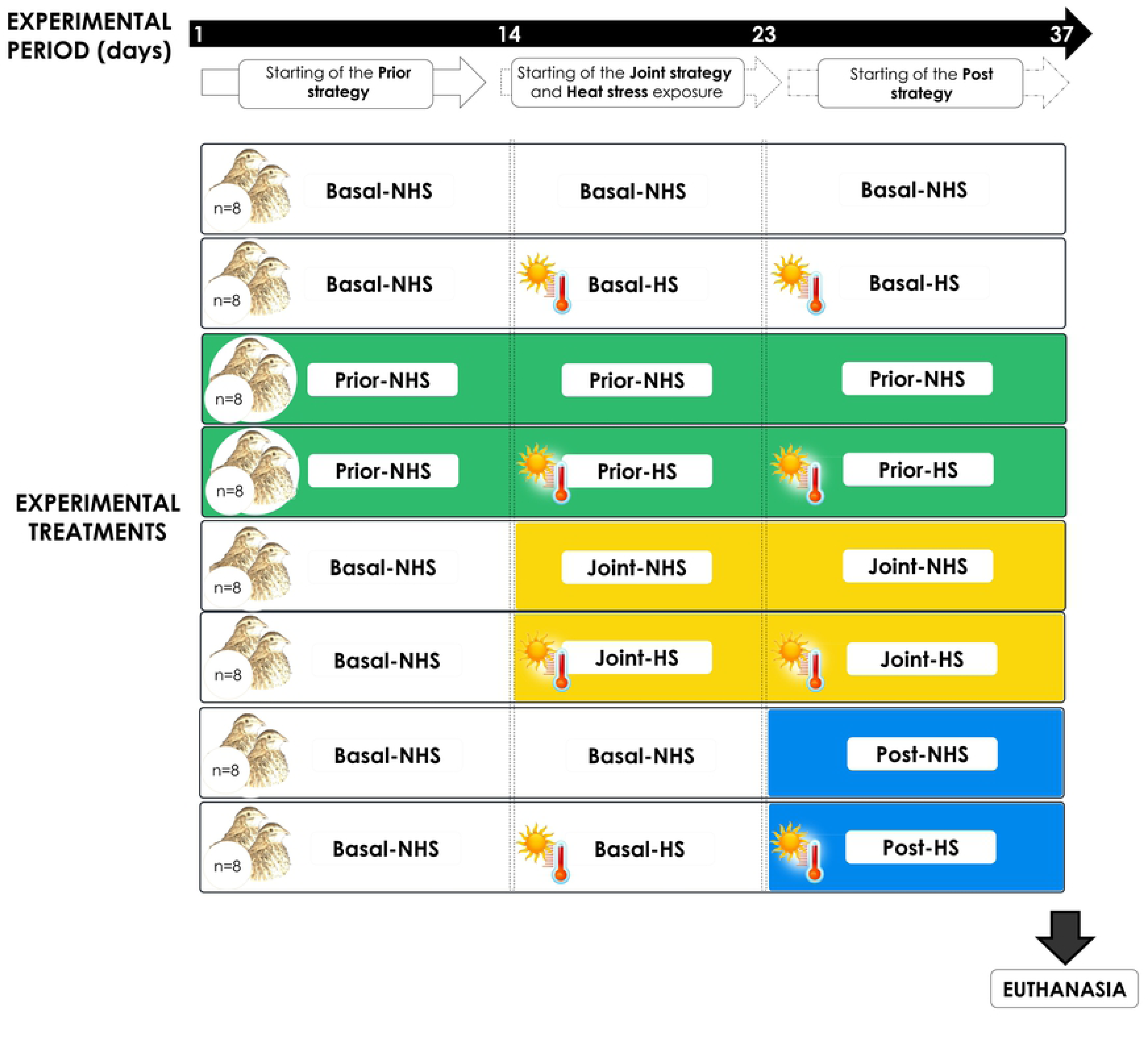
Timeline scheme. Days of the experimental period are indicated in the top horizontal arrow. All birds received a basal (Basal) diet and were maintained at standard environmental (room) temperatures (23.6 ± 0.1°C; NHS) until the experimental period began. From the first day of the experimental period and up to the end of it, thymol dietary supplementation began for females assigned to the Prior strategy (Prior). The remaining birds continued to receive the basal diet. The 14^th^ day of the experimental period, thymol dietary supplementation began for females assigned to the Joint strategy (Joint) and continued until the end of the experimental period. Simultaneously, heat stress (HS) was applied to half of the birds within each dietary group. From the 23^rd^ day of the experimental period (i.e., 9 days after HS was begun), thymol dietary supplementation started for females assigned to the Post strategy (Post), half of which had already been submitted to HS and the other half had continued at standard temperature. Finally, the 37^th^ day of the experimental period, all females were euthanized for biochemical analysis. Vertical dotted lines indicate application of a factor (i.e., diet or environmental temperature) for the corresponding experimental groups. A sample size of 8 birds was used for each experimental treatment.

### Assessment of physiological and behavioral traits

Given the potential sensibility of body-weight and feed intake as indicators of the transition between acute and chronic responses to the factors studied herein [21, 45], these traits were recorded throughout the experimental period (Table 1, and S3-S4 Appendix), to monitor the temporal evolution of the system. At the end of the experimental period, additional physiological and behavioral traits were assessed (Table 1 and S3, S4 and S5 Appendices). Specifically behavioral and egg production traits were registered between days 36 and 37 of the experimental period, and serum and liver samples were collected on day 37.

**Table 1.** Description of physiological and behavioral traits measured.

| Spatio-temporal scale | Subsystem | Trait | Description |
| --- | --- | --- | --- |
| Whole-organism level | Female somatic maintenance<sup>1</sup> | Body weight change | **Dynamics of body weight change** was estimated as ((final weight – initial weight) / final weight)*100; where final weight corresponds to body weight (g) on a specific day of the experimental period (days 10, 18, 26 or 34) and initial weight refers to body weight on day 2 of the experimental period. **Total body weight change** over the whole experimental period was estimated as ((final weight – baseline) / final weight)*100; where final weight corresponds to body weight (g) on day 34 and baseline to body weight on the week prior to initiation of experimental period (day -6). |
|  |  | Feed intake change | Feed intake (g) was estimated as the difference between the amount of feed supplied to each bird and the rest that remained in the feeders after two days and then accumulated over the same 8-day period used for body weight measurement. **Dynamics of feed intake change** was estimated as the difference in accumulated feed intake between a specific day (i.e., 10, 18, 26 or 34) and day 2 of the experimental period. **Total feed intake change** was estimated as the difference in accumulated feed intake between day 34 and baseline value corresponding to the week prior to initiation of experimental period (day -6). |
|  |  | Body temperature | Cloacal temperature (°C) measured on day 35 of experimental period at 9 h (morning) and 16 h (afternoon). |
|  |  | Respiratory rate | Number of thoracic movements per minute (bpm) measured on day 35 of experimental period at 9 h (morning) and 16 h (afternoon). |
|  |  | Panting | Average time spent panting per bird (s) measured on day 36 of experimental period, at 9 h (morning) and 14h (afternoon). |
|  |  | Droppings | The weight of droppings (g) deposited between 8-17 h measured on day 36 of experimental period. |
|  | Egg production<sup>2</sup> | Laying rate | Egg laying was registered daily and laying rate was estimated for the period between days 2 and 37 of experimental period. |
|  |  | Egg weight | Average weight of eggs collected on days 36-37 of experimental period. |
|  |  | Egg shell | Average percentage of the weight of the egg corresponding to the egg shell, of eggs collected on days 36-37 of experimental period. |
|  |  | Albumen | Average percentage of the weight of the egg corresponding to the egg albumen, of eggs collected on days 36-37 of experimental period. |
|  |  | Yolk | Average percentage of the weight of the egg corresponding to the egg yolk, of eggs collected on days 36-37 of experimental period. |
|  |  | Egg shape index | Average value of ((egg width / egg length) x 100) measured in eggs collected on days 36-37 of experimental period. |
| Molecular level | Serum and Liver<sup>3</sup> | TBARs | Lipid peroxidation indicator thiobarbituric acid reactive substances (nmol TBARS/mg protein) |
|  |  | Catalase | Activity of catalase enzyme (nano katal/ mg protein) |
|  |  | GPx | Activity of glutathione peroxidase enzyme (nano katal/mg protein) |
|  |  | FRAP | Estimation of reducing power with the ferric reducing ability assay (mM of Trolox equivalent/ g of liver or µL of serum) |
|  |  | TEAC | Estimation of antiradical activity with the trolox equivalent antioxidant capacity assay (mM of Trolox equivalent/g of liver or µL of serum). |
|  |  | TPC | Total polyphenol content (TPC) measured by Folin–Ciocalteu method (µg of gallic acid equivalents (GAE) / g of liver or µL of serum). |

For detailed description of methodology associated to each trait: <sup>1</sup>Female somatic maintenance (S3 Appendix), <sup>2</sup>Egg production (S4 Appendix), <sup>3</sup>Serum and Liver (S5 Appendix)

### Statistical analysis

Univariate statistical analyses were performed separately for each supplementation strategy. Thus, depending on the variable, a 2 x 2 x 3 or a 2 x 2 design that evaluated the effect of the diet supplied (basal and thymol), environmental temperature (NHS and HS) and day of experimental period (days 10, 26 and 34 in the Prior strategy and days 18, 26 and 34 in the Joint and Post strategies) or only the two first factors, respectively, was considered. We calculated a sample size of 8 female taking into consideration the experimental design and based on previously published data on the variability of body weight in laboratory quail populations [63]. Specifically, sample size was estimated using a standard variance of 16g [63], a minimum significant difference of 9g (3.5 % of body weight), a two-sided significance of 0.05 and a power of 0.8. Due to pair housing of females, feed intake and egg production could only be assessed at the cage level and not on an individual basis (n=4). No birds were considered outliers.

General Linear Mixed Models with the diet supplied, environmental temperature and day of the experimental period as fixed effects and wing band or cage number as random effects were used for the dynamics of body weight and feed intake, respectively. Detailed description of linear models and effects found for all 27 traits measured at the end-point of the experimental period can be found in S6 Appendix (Supplementary Tables S1-S4 and Figures S1-S4). Normality and homoscedasticity were evaluated both graphically (using tools of R library fitdistrplus) and numerically through Shapiro-Wilk, Levene’s and Bartlett tests. A *P*-value ≤ 0.05 was considered for statistical significance. LSD post hoc analysis was used to evaluate differences between treatments.

Multivariate statistics, clustering and Pearson correlations, were used to explore and describe overall pattern of variation in data and functional relationships among traits measured at the end-point of the experiment. Different clustering analyses were performed as follows:

1. An agglomerative hierarchical clustering analysis of the mean values obtained for each of the 8 experimental treatments was performed separately for each subsystem (female somatic maintenance, egg production performance, liver and serum antioxidant systems).
2. A “global” agglomerative hierarchical clustering analysis of the mean values of all traits obtained for each of the 8 experimental treatments. Since results of clustering on mean values could potentially be affected by individual variability, an agglomerative hierarchical clustering analysis using the individual values of all traits obtained from each female was also performed, which demonstrates the same overall pattern of means’ clustering (see details in S7 Appendix Fig S5).

In all clustering analyses, traits were first unit-based normalized (X′ = X − X_min_/X_max_ − X_min,_ where X is the set of observed values of a given trait, X_min_ and X_max_ are the minimum and maximum values present in X, respectively) thus, all values are between [0, 1]. Afterwards, Euclidean distance was calculated between traits and experimental treatments. With the corresponding similarity matrices, complete-linkage (the distance between two clusters is the longest distance between two points in each cluster) clustering method was used. Clustering was depicted in a dendrogram with a hierarchically-clustered heatmap.

Two main correlation analyses were performed as follows:

1. Pair-wise Pearson correlations were estimated between all traits independently of experimental treatment. These correlation coefficients and p-values were plotted in a heatmap when P<0.05.
2. Pair-wise Pearson correlations between all traits within each experimental treatment were estimated and represented as edges in a network when P<0.1 (see below network analysis). Further, the proportion of positive and negative correlations between subsystems for each experimental treatment was depicted as insets accompanying network representation (see below summary subsystems networks).

### Network analysis

#### Network definition

For each experimental treatment, we assumed a graph *G=*{**N**, **E, W**}, where **N** is the set of nodes (27 physiological and behavioral traits), **E** is the set of links (edges between de the nodes) and **W** denotes the edges’ weights, as a suitable model of a network [64]. We used a weighted adjacency matrix W_NxN_ to denote the edges’ weights, where an entry w_ij_ is a real number between [0–1] and represents the connectivity strength (absolute value of a Pearson correlation coefficient when P < 0.10) between the two nodes i, j, 0 < j < i < N [64] (*w_ij_* > 0 if a significant correlation exists between nodes *i* and *j*, otherwise, *w_ij_* = 0). Thus, the location of each nonzero entry in W specifies an edge for the graph, and the weight of the edge is equal to the value of the entry. For example, if W(2,1) = 0.90, then G contains an edge between node 2 and node 1 with a weight of 0.90, meaning that traits 2 and 1 are significantly correlated with a Pearson coefficient of 0.90.

The node degree *ki* is calculated by *ki* = Σ*j wij*, therefore, 0 ≤ *ki* ≤ *N* − 1 [65], and indicates the number of traits with which a given trait has significant correlations. Mean connectivity of the network was calculated as the average number of edges per node [66, 67].

#### Network entropy

For each of the 8 networks, Shannon’s entropy (H) was estimated as a measure of the uncertainty of a given variable [68]. Network entropy provides a robust measurement of the complexity of a weighted graph. It can be used as a descriptive statistic to compare the complexities between networks [64] that, for example, were constructed for two different experimental treatments. Specifically, we estimated the entropy associated with edges’ weight.

#### Edge entropy

Given a probability distribution p(x), where x is the probability of the weight of each edge, this entropy is defined as:

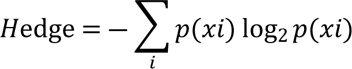

We estimated H_edge_, where p(x) is the probability that two traits are correlated with a given weight /correlation coefficient.

### Summary subsystems networks

For each experimental treatment, we assumed a graph *G=*{**N**, **E, W**}, where in this case **N** is the set of subsystems (female somatic maintenance, egg production, liver and serum antioxidant system) and each entry w_ij_ is given by the proportion of positive or negative correlations between two nodes. For example, W(2,1) = 0.11, indicates that G contains an edge between node 2 and node 1 with a weight of 0.11, meaning that subsystems 2 and 1 are significatively correlated in a proportion of 0.11 traits. The proportion of positive and negative correlations between subsystem within each experimental treatment was calculated as [C^(+/-)^**/**T], where C is the number of significant (P<0.1) positive (+) or negative (-) correlations detected between traits belonging to two given different subsystems and T the total number of possible correlations between all traits belonging to these subsystems. The total number of possible correlations between two given different subsystems T is [n_1_ * n_2_], where n_1_ and n_2_ are the number of traits belonging to each of these two subsystems.

### Software

Clustering analysis was performed using ScyPy and Seaborn methods (dendrogram and clustermap) in Python. MATLAB (RMatlab2018a) graph function was used for network construction. Custom MATLAB code was created for extracting network characteristics such as entropy and connectivity measures. Python and MATLAB codes are available in Figshare: https://figshare.com/s/d13f182242487f522376. All univariate analyses were performed with ‘R’ (The R Foundation for Statistical Computing) through a user-friendly interface implemented in InfoStat [69].

## Results

### Context- and time-dependency: divergent evolution of the system within each strategy is reflected in the changes in body weight and feed intake

The changes in body weight and feed intake over time showed a different pattern depending on the supplementation strategy (Figs 3 and 4, respectively).

**Fig 3.**
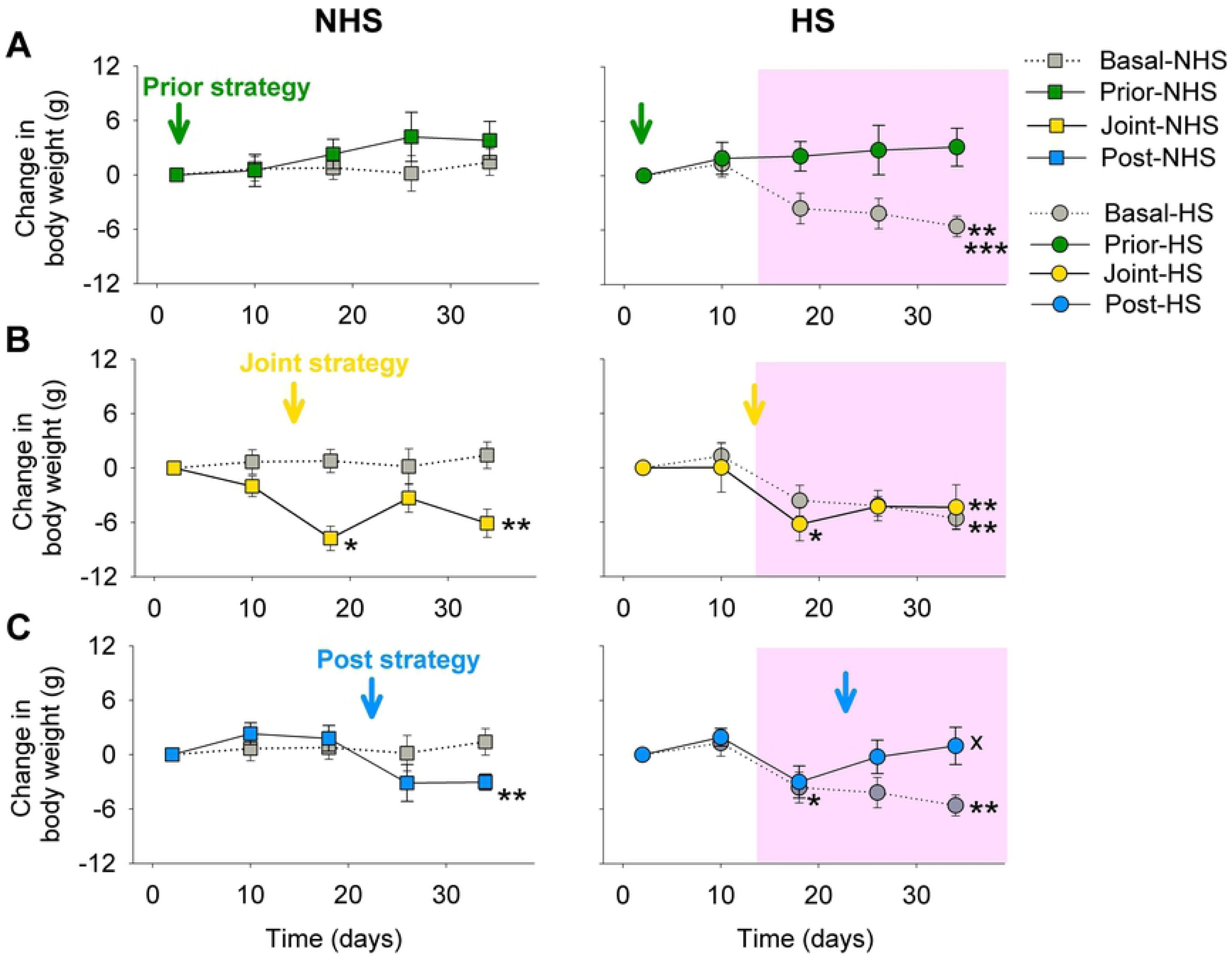
Dynamics of the effects of thymol dietary supplementation and chronic cyclic heat stress on body weight within each strategy. Mean (± sem) of the change in body weight was plotted as a function of days of the experimental period in the (A) Prior strategy, (B) Joint strategy and (C) Post strategy (n=8). Colored arrows mark the beginning of feed supplementation with thymol and the light red area indicates the beginning of the heat stress protocol (HS, circles), in the corresponding experimental treatments. Control birds not subjected to heat stress (NHS) are shown in squares, and those maintained on a basal diet in grey. To improve visualization, NHS and HS birds are shown separately on left and right panels, respectively. General Linear Mixed Models were performed on change in body weight at only three time points associated with initial, middle and end of experimental treatment, corresponding to days 10, 26 and 34 in the Prior and 18, 26 and 34 days of experimental period in the Joint and Post strategies. All the time points were depicted for reference. ***Indicate significant difference (P≤0.05) between Basal- HS and all other experimental treatments, independently of time point and temperature. **Indicate significant difference (P≤0.05) between experimental treatment and Basal-NHS, independently of time point. *Thymol supplementation significantly (P≤0.05) reduced body weight after 4 days of exposition. ^X^Indicates significant difference (P≤0.05) between thymol supplementation and Basal-HS at 34 d.

**Fig 4.**
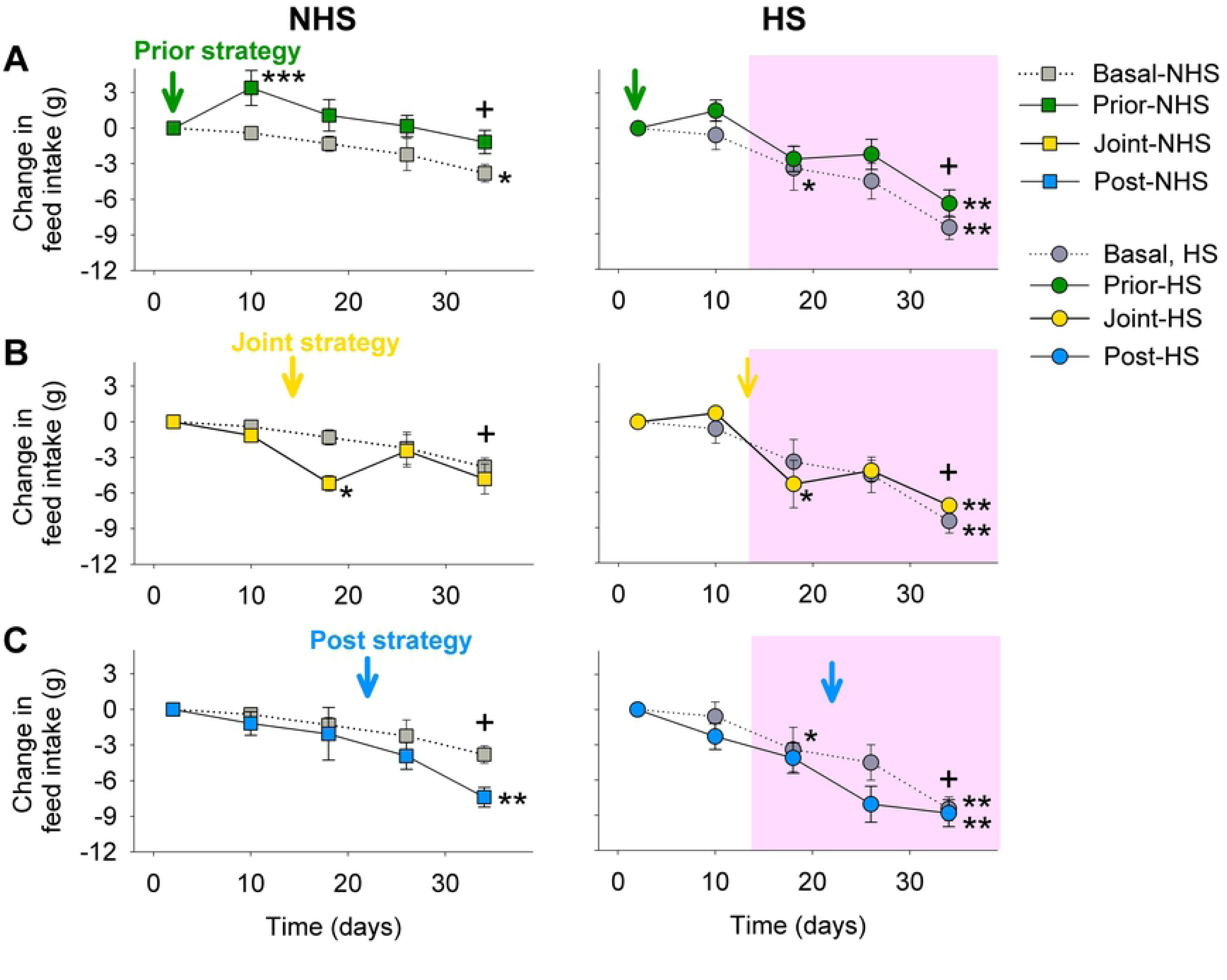
Dynamics of the effects of thymol dietary supplementation and chronic cyclic heat stress on feed intake within each strategy. Mean (± sem) of change in feed intake as a function of days of the experimental period in the (A) Prior strategy, (B) Joint strategy and (C) Post strategy (n=4). Colored arrows mark the beginning of feed supplementation with thymol and the light red area indicates the beginning of the heat stress protocol (HS, closed circles), in the corresponding experimental treatments. To improve visualization, birds not exposed to heat stress (NHS) and HS birds are shown separately on left and right panels, respectively. General Linear Mixed Models were performed on change in feed intake at only three time points associated with initial, middle and end of experimental treatment, corresponding to. days 10, 26 and 34 in the Prior and 18, 26 and 34 days of experimental period in the Joint and Post strategies. All the time points were depicted for reference. ***Thymol increased feed intake after 10 days. **Indicate significant difference (P≤0.05) between experimental treatment and Basal-NHS, independently of time point. *Thymol supplementation significantly (P≤0.05) reduced feed intake after 4 days. +Feed intake was lower (P≤0.05) on day 34 in comparison to days 18 and 26.

First, for body weight change in the Prior Strategy (Fig 3A), the diet in itself and in interaction with day of experimentation as well as the interaction between temperature and day of experimentation showed significant effects (Table 2), indicating that, under HS, thymol prevented the reduction in body weight observed in Basal-HS by days 26 and 34 of the experimental period (Fig 3A). Second, in the Joint strategy (Fig 3B), an effect of diet and significant interactions between diet and environmental temperature as well as diet and day were observed in the change in body weight (Fig 3B; Table 2). On day 18, thymol supplementation significantly decreased body weight compared to the Basal-NHS (Fig 3B). By the end of the experimental period, a lower body weight in all experimental treatments (Basal-HS, Joint-HS and Joint-NHS) was observed in comparison to control Basal-NHS. Lastly, in the Post strategy (Fig 3C), significant effects of interactions between diet and environmental temperature as well as diet and day of experimental period were observed in the change in body weight (Fig 3C; Table 2). By the end of the experimental period (day 34), under standard temperature, Post-NHS showed lower body weight values in comparison to Basal-NHS. However, under HS the opposite was observed and, by the end of the experiment, Post-HS had regained weight to values statistically similar to Basal-NHS birds (Fig 3C).

**Table 2.** Summary of statistical information regarding the effects of the diet supplied, environmental temperature, day of experimental period, and their interactions on the change in body weight and feed intake of adult female Japanese quail within each strategy.

|  | Change in body weight |  | Change in feed intake |  |
| --- | --- | --- | --- | --- |
|  | F-value | p-value | F-value | p-value |
| ***Prior Strategy*** |  |  |  |  |
| Diet | $F_{1,28}=5.31$ | **0,03** | $F_{1,12}=6.73$ | **0,02** |
| Envir. Temp | $F_{1,28}=1.32$ | 0,26 | $F_{1,12}=8.02$ | **0,02** |
| Day of exp period | $F_{2,56}=0.09$ | 0,91 | $F_{2,24}=73.59$ | **<0,001** |
| Diet x Environ. Temp | $F_{1,28}=1.06$ | 0,31 | $F_{1,12}=0.17$ | 0,69 |
| Diet x Day | $F_{2,56}=4.67$ | **0,01** | $F_{2,24}=0.25$ | 0,78 |
| Envir. temp x Day | $F_{2,56}=3.26$ | **0,05** | $F_{2,24}=8.36$ | **0,002** |
| Diet x Envir. Temp x Day | $F_{2,56}=0.96$ | 0,39 | $F_{2,24}=0.35$ | 0,71 |
| ***Joint Strategy*** |  |  |  |  |
| Diet | $F_{1,28}=7.26$ | **0,01** | $F_{1,12}=0.67$ | 0,43 |
| Envir. Temp | $F_{1,28}=2.95$ | 0,10 | $F_{1,12}=4.08$ | 0,07 |
| Day of exp period | $F_{2,56}=1.18$ | 0,31 | $F_{2,24}=15.45$ | **<0,001** |
| Diet x Environ. Temp | $F_{1,28}=5.37$ | **0,03** | $F_{1,12}=0.56$ | 0,47 |
| Diet x Day | $F_{2,56}=2.53$ | **0,05** | $F_{2,24}=5.45$ | **0,01** |
| Envir. temp x Day | $F_{2,56}=0.33$ | 0,72 | $F_{2,24}=2.64$ | 0,09 |
| Diet x Envir. Temp x Day | $F_{2,56}=1.21$ | 0,31 | $F_{2,24}=0.41$ | 0,67 |
| ***Post Strategy*** |  |  |  |  |
| Diet | $F_{1,28}=2.42$ | 0,13 | $F_{1,12}=2.03$ | 0,18 |
| Envir. Temp | $F_{1,28}=0.64$ | 0,43 | $F_{1,12}=4.96$ | **0,05** |
| Day of exp period | $F_{2,56}=2.10$ | 0,20 | $F_{2,24}=48.87$ | **<0,001** |
| Diet x Environ. Temp | $F_{1,28}=10.95$ | **0,002** | $F_{1,12}=0.03$ | 0,86 |
| Diet x Day | $F_{2,56}=3.22$ | **0,05** | $F_{2,24}=2.30$ | 0,12 |
| Envir. temp x Day | $F_{2,56}=0.11$ | 0,90 | $F_{2,24}=0.94$ | 0,40 |
| Diet x Envir. Temp x Day | $F_{2,56}=0.77$ | 0,47 | $F_{2,24}=0.17$ | **0,03** |

Change in body weight (n=8) and change in accumulated feed intake (n=4) was estimated as the difference between the measured value for each of the three time points (i.e., days 10, 26 and 34 in the Prior strategy; days 18, 26 and 34 in the Joint and Post strategies) and the value measured the day two after experimental period began.

Regarding change in feed intake, in the Prior strategy (Fig 4A), significant effects of all three factors as well as an interaction between temperature and day of experimental period were observed (Table 2). In general, feed intake decreased over time except in Prior-NHS that even showed an increase by day 10 in comparison to Basal-NHS. A similar overall pattern was observed in the Joint and Post strategies, but with a key difference. Contrary to that observed in the Prior strategy, both the Joint-NHS and the Post-NHS decreased feed intake on the first days following initiation of supplementation compared to Basal-NHS (Fig 4B and 4C; Table 2). Moreover, in the Post-NHS, at the end of the experimental period, feed intake continued to be lower than Basal-NHS.

Note that each supplementation strategy underwent distinct dynamical changes and key effects of the respective temporal evolutions are still observable by the end of the experimental period.

### Functional relationships among traits within each subsystem differ among experimental treatments

The role of the context on the functional relationships among traits within each subsystem is shown in Figure 5. In each panel, the x-axis presents the traits while the y-axis presents experimental treatments. The corresponding dendrograms and heatmaps describe the similarity among the clusters found, and show distinct grouping of the experimental treatments depending on the subsystem considered (Fig 5).

**Fig 5.**
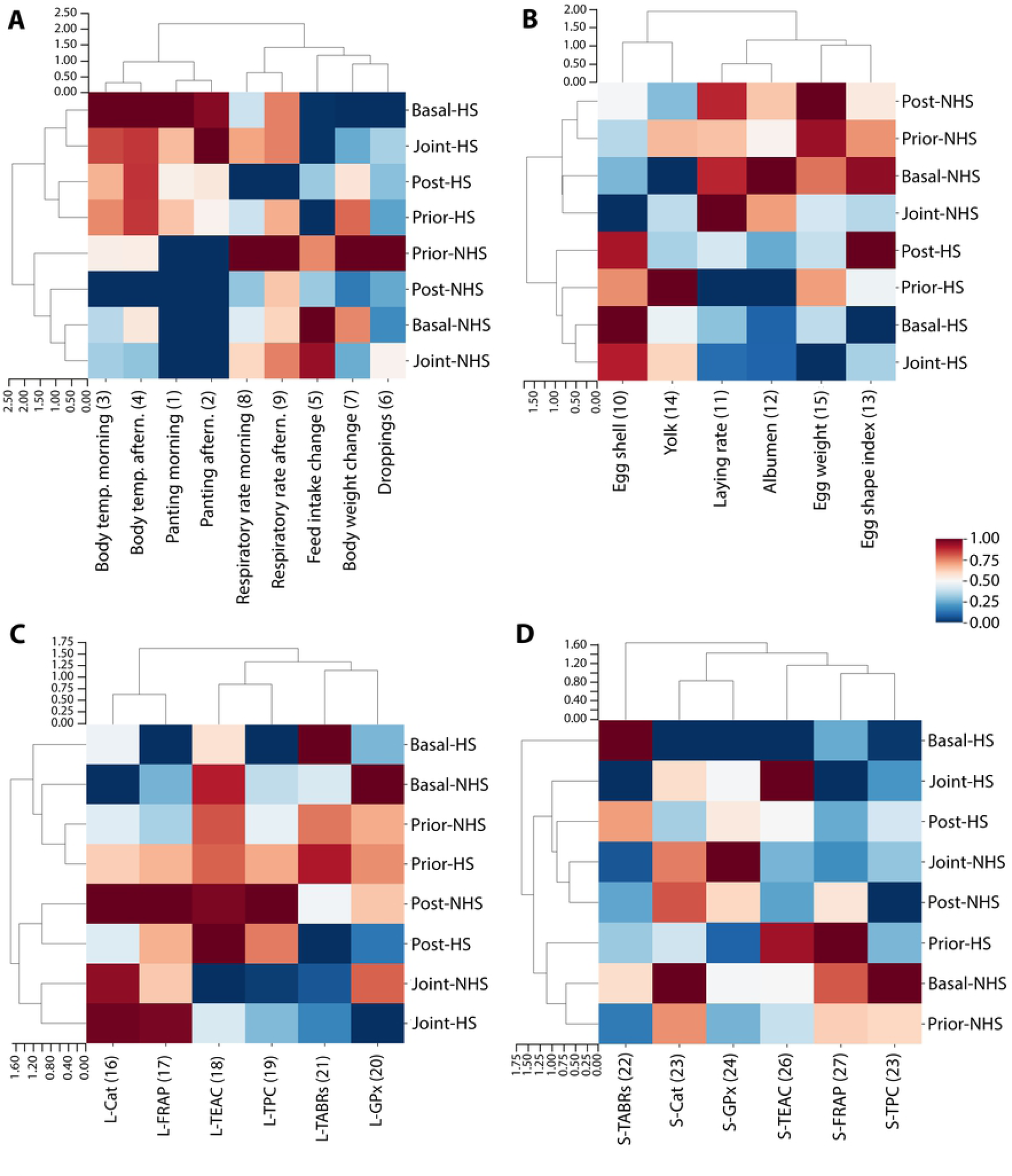
The functional relationship among physiological and/or behavioral traits within each subsystem. Agglomerative hierarchical clustering (complete-linkage method) and heatmap of the mean values obtained in each treatment for traits belonging to each subsystem, namely (A) female somatic maintenance, (B) egg production, and (C) liver and (D) serum antioxidant response of adult female Japanese quail. Experimental treatments are described in Fig 2. Heatmap shows from dark blue to dark red the lowest to highest normalized values of the traits. Clustering of experimental treatments and traits are shown in y- and x- axis, respectively. Prefixes “L-” and “S-” in traits’ names denote they correspond to liver and serum, respectively. TBARs= thiobarbituric acid reactive substances; TEAC= trolox equivalent antioxidant capacity; FRAP= ferric reducing ability; TPC= total polyphenol content; Cat= activity of catalase enzyme; GPx= activity of glutathione peroxidase enzyme. Body weight change and feed intake change correspond to the total change in those traits (estimated as indicated in Table 1). Numbers in parenthesis after trait name are coincident with the ordering of traits in networks shown in Fig 7.

In female somatic maintenance (Fig 5A) and egg production (Fig 5B) subsystems, the two most distant clusters of experimental treatments observed in the y-axis dendrogram are directly associated with environmental temperature. This is principally explained by the significant increase in body temperature and panting (red region of heatmap in Fig 5A,) or the increase in the percentage of egg shell (red region of heatmap in Fig 5B), as well as decrease in laying rate and percentage of albumen (blue region in heatmap Fig 5B), under heat stress. For the corresponding univariate analysis see S6 Appendix Fig S1 and S2, as well as Tables S1 and S2. Within these main clusters, independently of environmental temperature, the Joint strategy is more similar to the Basal diet, than to the Prior and Post strategies (Fig 5A, B). Grouping of the Prior and Post strategies is supported by significant effects of the diet, including reduction of body temperature and time spent panting as well as increment of body weight and feed intake (S6 Appendix Fig S1, Table S1). Oppositely, grouping of the Basal diet and the Joint strategy is associated with the inability of thymol in the latter to promote changes in most of the traits and even reduce egg weight (compare with S6 Appendix Fig S2E, Table S2). As an exception, all three supplementation strategies avoided the reduction in the weight of droppings induced by HS (S6 Appendix Fig S1S-S1U, Table S1). In other words, in these subsystems, most of the variability in the response among experimental treatments is given by the environmental temperature, although some specific and more subtle context-dependent effects of the diet are observed.

Contrary to that observed in the two previous subsystems representative of the whole-organism scale, in the liver antioxidant system response, the most distant clusters in the y-axis dendrogram, group the experimental treatments according to diet (Fig 5C, y-axis). On one side, the Basal and Prior strategy are grouped together independently of environmental temperature, Prior- NHS and Prior-HS being more similar to Basal-NHS than to Basal-HS. Note that, with the exception of catalase activity, univariate analyses in the Prior strategy did not detect significant effects of the diet or the temperature for none of these traits (S6 Appendix Fig S3 and Table S3). On the other side, the Joint and Post strategies are grouped together independently of environmental temperature, and specific effects of the diet can be observed (see also univariate analysis in S6 Appendix Fig S3 and Table S3). Grouping of the Joint and Post strategies is primarily supported by catalase activity which more than doubled its value in the Joint-NHS and Post-NHS compared to the Basal-NHS (S6 Appendix Fig S3E and S3F) and increment in the reducing power observed in the Joint and Post strategy in comparison to the Basal diet (L-FRAP, S6 Appendix Fig S3K and S3L). Additionally, reduced lipid peroxidation was observed in the Joint-HS, Joint-NHS and Post-HS in comparison to their diet controls; while this reduction was not detected in the Prior strategy (L-TBARS, S6 Appendix Fig S3A-C). However, in the Joint strategy, it is also noticeable the generally lower values of all traits (Fig 5C blue color in heatmap) and the significant effect of both factors diet and environmental temperature in lipid peroxidation (L-TBARS, S6 Appendix Fig S3B). These results highlight the importance of context-specific thymol regulation of the liveŕs antioxidant response.

The serum subsystem is unique (Fig 5D) given that in the y-axis dendrogram the Basal-HS appears alone, separated from the clusters containing all the other experimental treatments (Fig 5D, y-axis). Basal-HS shows comparably higher values (red in heatmap) of lipid peroxidation (S-TBARs) and lower values (blue in heatmap) of the other traits, in comparison to the rest of the experimental treatments (compare with S6 Appendix Fig S4, Table S4). The grouping of the Prior-HS with Prior- NHS and Basal-NHS treatments denotes the ability of the Prior strategy to prevent the effect of heat stress in the serum antioxidant system response. This effect is most notable in antioxidant capacity measured as reducing power (S-FRAP, S6 Appendix Fig S4J, Table S4) and antiradical activity (S- TEAC; S6 Appendix Fig S4M, Table S4) denoted as red regions of the heatmap, although significant effects are also observed in serum catalase activity (S-Cat; S6 Appendix Fig S4D, Table S4). In regard to the Joint and Post strategies, although they presented a similar response pattern, specific univariate statistical differences were detected, reflecting differential magnitude of effects. For example, significantly higher values (Fig 5D, red regions of the heatmap) of serum catalase activity and antiradical activity (S-TEAC) are observed in Joint-HS in comparison to the HS control, which was not observed in Post-HS (see also S6 Appendix Fig S4E and S4N, Table S4). Thus, although modulation on the upon serum antioxidant system components under heat stress are observable in all supplementation strategies, heat stress impacts to a lesser extent in the Prior strategy.

### Overall pattern of variation and relationships among whole-organism and molecular-scale traits

The role of context in the general response pattern integrated over the different subsystems is shown in Figure 6A.

**Fig 6.**
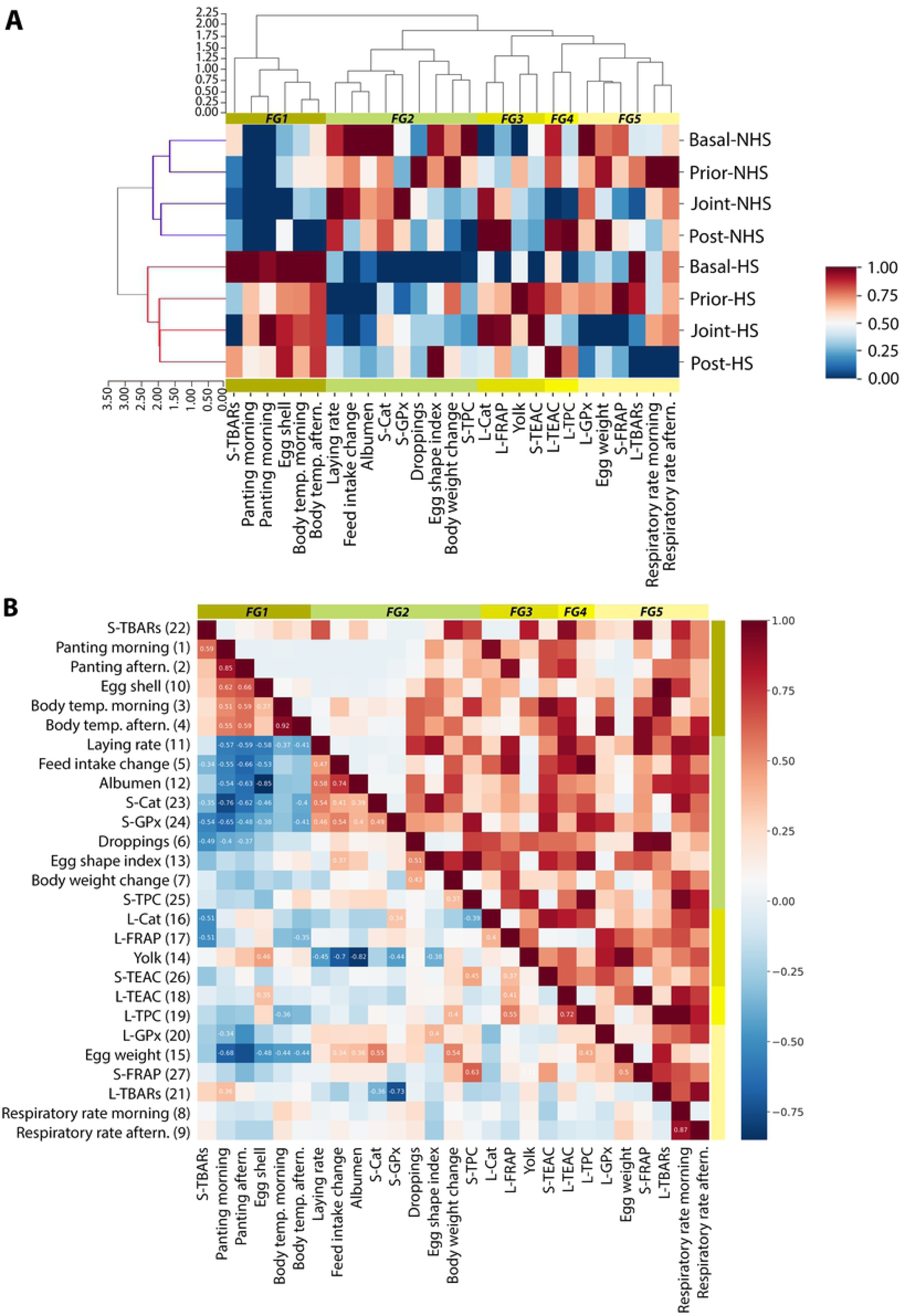
Functional relationships among all physiological and behavioral traits and emergent effects of thymol supplementation and heat stress. (A) Agglomerative hierarchical clustering (complete-linkage method) and heatmap of the mean values obtained in each treatment for all traits measured in adult female Japanese quail. Experimental treatments are described in Fig 2. Heatmap shows a color scale from dark blue to dark red representing the lowest to highest normalized values of the traits. Clustering of experimental treatments and traits are shown in y- and x- axis, respectively. Numbered “FG” labels in the green-scaled horizontal bar indicate functional groups emergent from traits clustering. Prefixes “L-” and “S-” in traits namés denote they correspond to liver and serum, respectively. Numbers in parenthesis after trait name are coincident with the ordering of traits in networks shown in Fig 7. (B) Pearson correlation coefficient (lower triangle) and its corresponding p-value (upper triangle) between each pair of traits are represented, ordered according to Fig 6A. Specifically, the color scale from dark blue to dark red indicate the range from the lowest to highest correlation coefficients (-1 to 1). Coefficient values of significant correlations (P < 0.1) are annotated in the lower triangle. Numbered “FG” labels in the green-scaled horizontal bar indicate functional groups emergent from traits clustering in Fig 6A.

The most distant clusters of experimental treatments in the y-axis dendrogram denote a grouping based on environmental temperature (6A, y-axis). Within the NHS cluster, 2 more clusters emerge associated with diet, grouping the Prior strategy with the Basal diet on one side, and the Joint with the Post strategy on the other. Within the HS cluster, the three thymol supplementation strategies were grouped together and separate from the Basal diet, highlighting that, globally, all three strategies produced physiological and behavioral changes in comparison to the Basal diet.

In relation to traits’ clustering, 5 functional groups emerge (FG, Fig 6A green-scaled colored rectangles along x-axis). Interestingly, the distribution of the traits in the functional groups is non-trivial, given that they do not mark division between subsystems. Instead, functional groups reflect how the modulation of each trait in particular emerges from the complex interplay between processes occurring at different spatio-temporal scales and subsystems; resulting in the relative increase (Fig 6A, red in heatmap) or decrease (Fig 6A, blue in heatmap) of the trait’s value.

The first two functional groups (Fig 6A, FG1 and FG2) are mostly associated with general changes induced by HS (see details of each trait in previous sections), comprising traits from the female somatic maintenance, egg production, and serum subsystems. Notably, while traits in functional group 1 increase, those in functional group 2 decrease under HS. Since the traits from each of these functional groups respond to HS in an opposite fashion, the traits within each functional group are positively correlated with each other while they are negatively correlated with the traits from the other functional group (Fig 6B). Thus, potentially denoting the coordinated functional response to HS of the traits within these two functional groups.

The following 3 functional groups (Fig 6A, x-axis dendrogram) are principally associated with the context-specific thymol regulation of the serum’s and liveŕs antioxidant response described in the previous section (Fig 5C and 5D). Pattern of variation of the traits in these functional groups highlights thymol potential for improving antioxidant system through divergent modulation of its components. This treatment-specificity is reflected in the generally low number of significant correlations that exhibit these traits with other traits within the same or other functional groups (Fig 6B).

### Network analysis of the response to heat stress and thymol dietary supplementation

Networks characteristics vary substantially depending on the experimental treatment (i.e., context) the birds are subject to (Fig 7 and Table 3).

**Fig 7.**
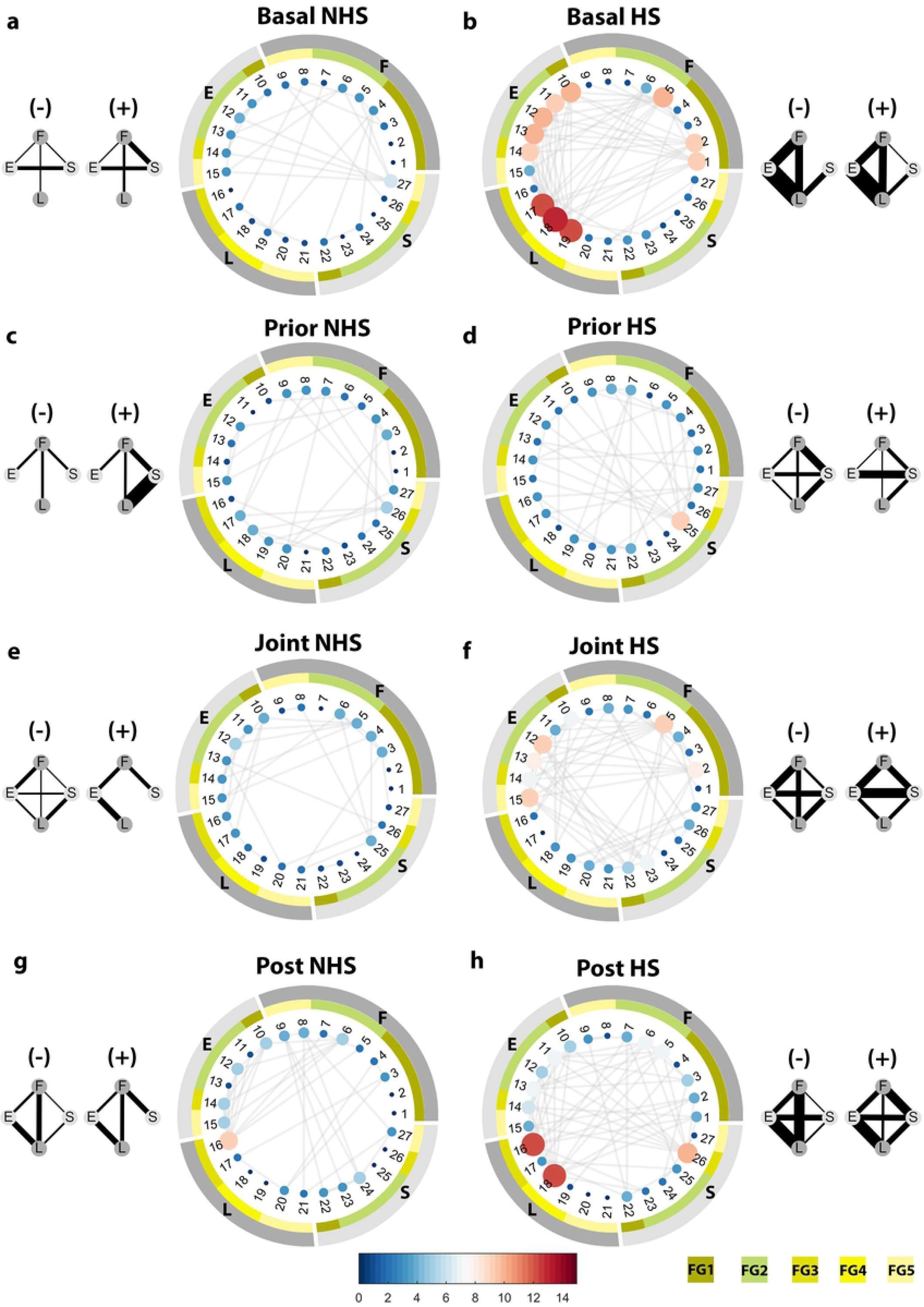
Networks of physiological and behavioral traits across spatio-temporal scales and subsystems. Pearson correlations (when *P*<0.1) among traits of adult female Japanese quail from the basal diet under (A) standard temperature (Basal-NHS) and (B) heat stress (Basal-HS), the Prior strategy under (C) standard temperature (Prior-NHS) and (D) heat stress (Prior-HS), the Joint strategy under (E) standard temperature (Joint-NHS) and (F) heat stress (Joint-HS), the Post strategy under (G) standard temperature (Post-NHS) and (H) under heat stress (Post-HS). Gray edges represent significant correlation between traits. Edges thickness is given by the value of the correlation coefficient. In each network, external gray circle identifies the subsystems the traits belong to and green-scale colored internal circle identifies functional groups (FG) obtained in Fig 6A. Nodes represent physiological or behavioral traits. Node size and color is given by its degree (i.e., number of significant correlations with other traits), from dark blue to dark red it shows the lowest to highest degrees. Node numbered label corresponds to the ordering of traits emergent from clustering in Fig 6A nested within each subsystem: 1. Panting morning, 2. Panting afternoon, 3. Body temp. morning, 4. Body temperature afternoon, 5. Feed intake change, 6. Droppings, 7. Body weight change, 8. Respiratory rate morning, 9. Respiratory rate afternoon, 10. Egg shell, 11. Laying rate, 12. Albumen, 13. Egg shape index, 14. Yolk, 15. Egg weight, 16. L-Cat, 17. L-FRAP, 18. L-TEAC, 19. L-TPC, 20. L-GPx, 21. L-TBARs, 22. S-TBARs, 23. S-Cat, 24. S-GPx, 25. S-TPC, 26. S-TEAC, 27. S-FRAP. Letters in summary subsystem networks (insets) indicate the different subsystems: F for female somatic maintenance; E for egg production; L for liver; S for serum.

**Table 3.** Descriptive measurements of the networks constructed from 27 physiological and behavioral traits of adult female Japanese quail fed on a basal diet or thymol supplemented within each of three strategies (Prior, Joint, Post), and subjected either to standard or heat stress environmental temperature.

| *Network characteristic* | **Basal** |  | **Prior Strategy** |  | **Joint Strategy** |  | **Post Strategy** |  |
| --- | --- | --- | --- | --- | --- | --- | --- | --- |
|  | NHS | HS | NHS | HS | NHS | HS | NHS | HS |
| Number of Nodes | 22 | 27 | 23 | 27 | 23 | 26 | 21 | 25 |
| Number of Edges | 26 | 73 | 30 | 37 | 31 | 58 | 37 | 60 |
| Average Weight of Edges | 0,98 | 0,97 | 0,98 | 0,99 | 0,98 | 0,98 | 0,99 | 0,99 |
| Edge's Weight Entropy | 1,14 | 2,47 | 0,75 | 1,08 | 1,21 | 2,00 | 0,84 | 1,31 |
| Mean Connectivity | 1,18 | 2,70 | 1,30 | 1,37 | 1,35 | 2,23 | 1,76 | 2,40 |

Supplementation started 14 days before, simultaneously or 9 days after the beginning of HS exposure in the Prior, Joint and Post strategy, respectively, and continued up to the end of the experimental period.

In the Basal-HS an increased level of connectivity and entropy (Table 3 and edges in Fig 7B) is observed in comparison to all other experimental treatments (Fig 7A and 7C-7H, Table 3). Noticeably, under heat stress, the nodes representing liver indicators of antioxidant capacity (Fig 7B, nodes 17, 20 and 21) are particularly influential, as they present heightened level of connectivity with traits of female somatic maintenance and egg production (Table 3 and Fig 7B, light pink nodes).

Concomitantly, the presence and proportion of correlations among subsystems is increased in Basal- HS in comparison to the standard temperature, the liver subsystem becoming particularly important in the communication with other subsystems under heat stress (insets Fig 7). Similarly, but to a lesser extent, within each supplementation strategy connectivity increases under heat stress in comparison to standard temperature (compare Fig 7C, E, G with Fig 7D, F, H and Table 3) although different nodes are affected. This increase is less pronounced in the Prior than in the Joint and Post strategies, while avoiding any node becoming predominant over others as observed by the highest values of entropy in this group (Table 3). Equivalently, almost all subsystems are correlated within each supplementation strategy under heat stress, being the Prior strategy that in which heat stress effects have a lesser extent as seen in the lower value of the proportion of correlations. Thus, the network’s connectivity and entropy properties as well as level of correlation among subsystems can be differentially modulated depending on the moment dietary supplementation begins in regard to the onset of heat stress.

## Discussion

Recognizing that biology is fundamentally contextual [70], is crucial to understand how the interaction of myriad components gives rise to the dynamic and complex behavior of biological systems [71]. Herein, supplementation strategies were shown to evolve differently over the 5-week experimental period in regard to body weight and feed-intake. Under standard environmental temperature, thymol supplemented females in the Prior strategy showed no change in body weight, while females in the Joint and Post strategies evidenced a reduction in that trait (Fig 3, respectively) in comparison to the Basal diet. Discrepancies among strategies are multifactorial, and may be attributable in part to the duration of the supplementation period applied in each one of them (i.e., approximately 5 weeks, 3 weeks, and 2 weeks in the Prior, Joint and Post strategies, respectively). A previous work in growing broilers has shown that initial reduction in body weight promoted by thymol can be recovered over time [72]. However, it is important to note that, in many studies, effects of thymol on the change in body weight are not always evident [63, 73] as observed in the Prior strategy or, on the contrary, weight gain can occur [42, 72]. Similarly, apparent contradictory effects of thymol have also been reported regarding feed intake. On one hand, the inclusion of thymol in the diet of pigs [74], poultry [15,75–78], and specifically female quail in the short-term quail [45] can promote reduced feed intake. On the other hand, several studies evaluating the long-term effects of thymol supplementation have shown that feed intake was unchanged after chronic thymol supplementation (>3-weeks) in growing broilers [42, 73], adult laying hens [44] and Japanese quail [63, 79], as observed in the Joint and Prior, but not in the Post, strategies. In a more detailed study, during the first 6-weeks of feed supplementation in growing broilers, mostly no significant effects on weekly feed intake were observed in comparison to controls [72]. Thus, in poultry, it seems that sensibility to thymol decreases over time, possibly due to a habituation process.

It is also important to take into account that thymol supplementation in the Prior strategy prevented and in the Post strategy partially reversed the expected heat-induced weight loss [21,80– 83], without increasing feed intake. Thus, sequential exposure to the novel stimulus (i.e., novel feed supplement or elevation of environmental temperature) could favor a certain level of adaptation to one of them before the exposure to the next potentially stressful stimulus, as reported for thymol supplemented animals previous or after exposition to a variety environmental challenges [54–56]. On the contrary, joint exposure (Joint strategy) to both novel and potentially stressful stimuli could overwhelm, at least temporarily, the capacity of the birds to cope with them, resulting in the inability to recover weight loss. Thus, by the end of the 5-week experimental period, birds from each experimental treatment had experienced a unique trajectory associated with either sequential or joint exposure to heat stress and dietary supplementation. These findings suggest that the outcome observed at the end-point of the experimental period should be conceived as a contingency or dynamical state of the transition between short- and long-term response to each of these two factors within a particular trajectory.

At the end-point of this experimental period, the divergent modulation of the subsystems by the supplementation strategy brought to light the differential sensibility of each subsystem to the factors applied. First, the observation that the response of female somatic maintenance and egg production subsystems is dominated by heat stress (Fig 5A,B) is logical given that many of the traits involved are known to be directly associated with thermoregulation and energetic balance [80, 84]. Also, the detrimental effects of heat stress observed on these traits are consistent with precedent reports for adult quail exposed to chronic cyclic heat stress [27,50,85–87]. Contrarily, the liver response was primarily associated with thymol effects (Fig 5C). This may be attributable to the role of the liver as xenobiotics’ detoxifying organ [88] as well as its fundamental role in lipid metabolism [89, 90]. In this context, several liver lipid metabolism pathways are known to be modulated by thymol [40]. Interestingly, liver TBARs is sensitive to both heat stress [14] and thymol[16,38,40], and behaves as a key outcome of the underlying interplay between these factors at the local level. Lastly, the serum subsystem mainly showed that thymol in all three supplementation strategies was able to modulate the response to heat stress (Fig 5D). Thus, this level seems to par excellence reflect the effects of both factors. This finding is reasonable considering that serum antioxidant system biomarkers result from both translocation of oxidation end-products from tissues (e.g., liver, muscle, heart, etc.) to systemic circulation and oxidative process in cells of peripheric circulation [91, 92]. Thus, on one hand, these biomarkers reflect the trade-off between pro-oxidant effects directly attributable to heat stress-induced mitochondrial dysfunction in tissues [14,29,93] and the strong opposite modulation of this process at the local level by thymol [38,40,94]. On the other hand, they also reflect sustained inflammation and global pathophysiological condition imposed by heat stress [14,91,92]. Hence, the differential sensibility to the factors inherent to each subsystem should be taken into account to understand the integrated-organismic response within each strategy.

In the broader framework of complex systems, self-organization and network physiology [7,95,96] an important question is how multi-component coordination among processes occurring at different spatio-temporal scales generates emergent behavior at the integrated level [97–100]. The approach applied herein demonstrates that the collective behavior of all interacting traits is non-trivial (Figs 5,6,7). Specifically, network analyses revealed that coordinated responses among traits (i.e., correlations) are context-specific (Fig 7). The high level of correlation observed among traits and subsystems under heat stress (Fig 7B) is consistent with this factor acting as the main constraint (Fig. 1) imposed on the system (i.e., defines initial and boundary conditions) [4, 101], with organism level responses (i.e. increased panting and body temperature as well as decreased body weight and feed intake) cascading downwards. In turn, the molecular response to heat stress could influence higher levels through dynamic changes [102] associated with the antioxidant response. Hence, heighted modulation in one trait under HS would also be observed in other traits from different subsystem within a given bird. This is most noticeable in Basal-HS, where highly correlated traits include L- FRAP, L-TEAC, L-TPC, Panting, Feed intake, Albumen, egg-shape index, and egg shell. Note that these first three molecular traits are not actual components of metabolic pathways but rather sensors of the liver’s redox environment. The high level of connectivity of these liver traits could be associated with previous studies that have proposed that the liver can act as a metabolic hub under certain conditions such as adaptation to temperature, being this organ key in the overall regulation of the balance between energy intake and expenditure [103, 104]. Moreover, results are consistent with the proposal that a redox network could have an important role in sharing information and eliciting responses to that information in a coordinated manner throughout the entire body [8]. In all, our observations support the contention that highly connected hubs play a disproportionate role either in influencing the expression patterns of other nodes in the network or alternatively acting as “sentries” communicating changes that occur elsewhere in the network [105].

The high level of connectivity observed in Basal-HS decreased in birds under heat stress and supplemented dietary thymol (compare Fig 7B,D,F,H). The modulation observed is consistent with the more general framework indicative of the antagonistic multiscale and scale-dependent effects of heat stress and thymol supplementation, influencing from gene expression and metabolic pathways to the whole-organism (see [14,18,27,46–51,19–26] for heat stress, [16,33,42–44,34–41] for thymol, and [21,45,53,54,62] for their combination). However, our experimental approach allowed us to further demonstrate that the resultant non-linear combination of constraints imposed on the system by both factors emerges as a more variable or “less coordinated” response pattern (i.e., lower level of correlations among traits and subsystems, lower entropy and connectivity, general absence of dominant nodes) in comparison to the Basal-HS. Moreover, the exact network configurations showed important variations among experimental treatments, as well as the level of positive and negative correlations among subsystems.

As we hypothesized, thymol potential for multi-scale modulation of heat stress consequences on female quail is dependent on the moment in which supplementation initiates with regard to the beginning of this environmental challenge. Changes observed indicate that thymol supplementation can prevent (Prior strategy), attenuate (Joint strategy) or even reverse (Post strategy) some of the effects of heat stress. In general, sequential exposition to the factors/stimuli resulted in beneficial modulation of traits. Particularly the Prior strategy showed the greater-extent modulation of heat-induced deleterious effects. The only deleterious effect of heat stress that couldn’t be modulated by thymol in this strategy was the increment of liver’s TBARs (Fig 5C also S6 Appendix Fig S3A). This observation can be explained by potential pro-oxidant and cytotoxic effects of thymol [106] resultant of the longer supplementation period within this strategy and also the increased excretion observed that may be conducive to marginal deficiencies of minerals and vitamins that act as direct antioxidants and/or cofactors of antioxidant enzymes [107–109]. Thymol within the Post strategy promoted enhanced thermoregulatory effects. It not only increased the weight of droppings, possibly reflecting an increase in water intake [54], as seen in the other two strategies, but also decreased body temperature and panting (Fig 5A also S6 Appendix Fig S1C,F,O,R,U). These findings can be associated with thymol’s cardiovascular effects [110–112] and several unspecific thermoregulatory effects [113–115] in a context of favored energetic balance, i.e., low feed intake and low body weight and, thus, lower metabolic heat production [74, 116].

In contrast, the Joint strategy not only didńt show some of the beneficial effects observed in the other two strategies, but even induced a detrimental decrease in egg weight both under heat stress and standard temperature (Fig 5B also S6 Appendix Fig S2E). These last aspect is opposite to previous studies of thymol supplementation that showed either no significant effects on egg weight nor changes in feed efficiency [63,79,117] or an increment in egg weight associated with an increase in feed efficiency [44]. In this manner, in the Joint strategy, the observed initial reduction of feed intake could have influenced nutrient availability for egg formation [118–120], as well as impair overall energetic balance. In all, these results, further highlight that different constraint and causation processes could have modulated the configuration of the network [101] (see Fig 1) in each strategy as a consequence of the distinctive trajectories experienced by them (Figs 3 and 4). Therefore, the combination of univariate and multivariate approaches such as the ones applied here, that consider traits as emergent properties of a complex system, can begin to shed light on previously counterintuitive and/or inconsistent results among different experimental settings [15–17].

Previous research studying cross-correlations or coherence between time series of different traits found a robust relation between network structure and physiologic states: every state is characterized by specific network topology, node connectivity and links strength [7]. It is important to note, that with our experimental design we were not able to perform these type of time series analyses. First, because serum sampling is invasive and can act as a stressor in itself, only one sample was taken. Second, and most important, because liver samples were obtained post-mortem. However, based on our results it is reasonable to assume that, similar to physiological states associated with sleep/wake states [6,121,122] or pathological conditions [123, 124], environmental challenges as well as dietary supplements affecting the antioxidant system have the potential to induce distinct, well defined, physiological states. Further studies that include analysis [1, 125] of a variety of biomarkers in live birds over time are needed to fully describe these particular physiological states.

## Conclusions

Accounting for their multiple-multi-scale and context-dependent roles *in vivo*, both together or separately, thymol dietary supplementation and chronic cyclic heat stress have the potential to alter somatic maintenance of female quail as well as local and systemic antioxidant response. By understanding the traits assessed as nodes of a physiological network, we were able to provide evidence of divergent functional relationships among them under the different experimental treatments studied. Our findings also highlight the differential roles of the serum and liver antioxidant systems, and the complex nonlinear association among molecular mechanisms of redox biology, resource allocation and metabolism. In all, the response pattern observed could be understood as a reflection of an emergent physiological state induced by the environmental challenge and dietary supplementation. Our work constitutes the first study that includes network, integrative, and experimentally comparative analyses applied to the field of dietary supplementation under environmental challenges. This perspective could help understand complex biological responses, which is important for developing efficient and welfare orientated supplementation protocols and interpreting the apparently paradoxical background in this field.

## Acknowledgments

We thank Maria Julia Ortiz and Pablo Prokopiuk for technical assistance with animal husbandry. We also acknowledge PhD. Vet. Martin Jorge Caliva, PhD. Biol. Franco Nicolas Nazar, Biol. Emiliano Ariel Videla, Biol. Octavio Giayetto, Biol. Cristian Jaime, Biol. Gabriel Orso, Biol. Florencia Córdoba, for technical assistance with collection of biological samples.

---

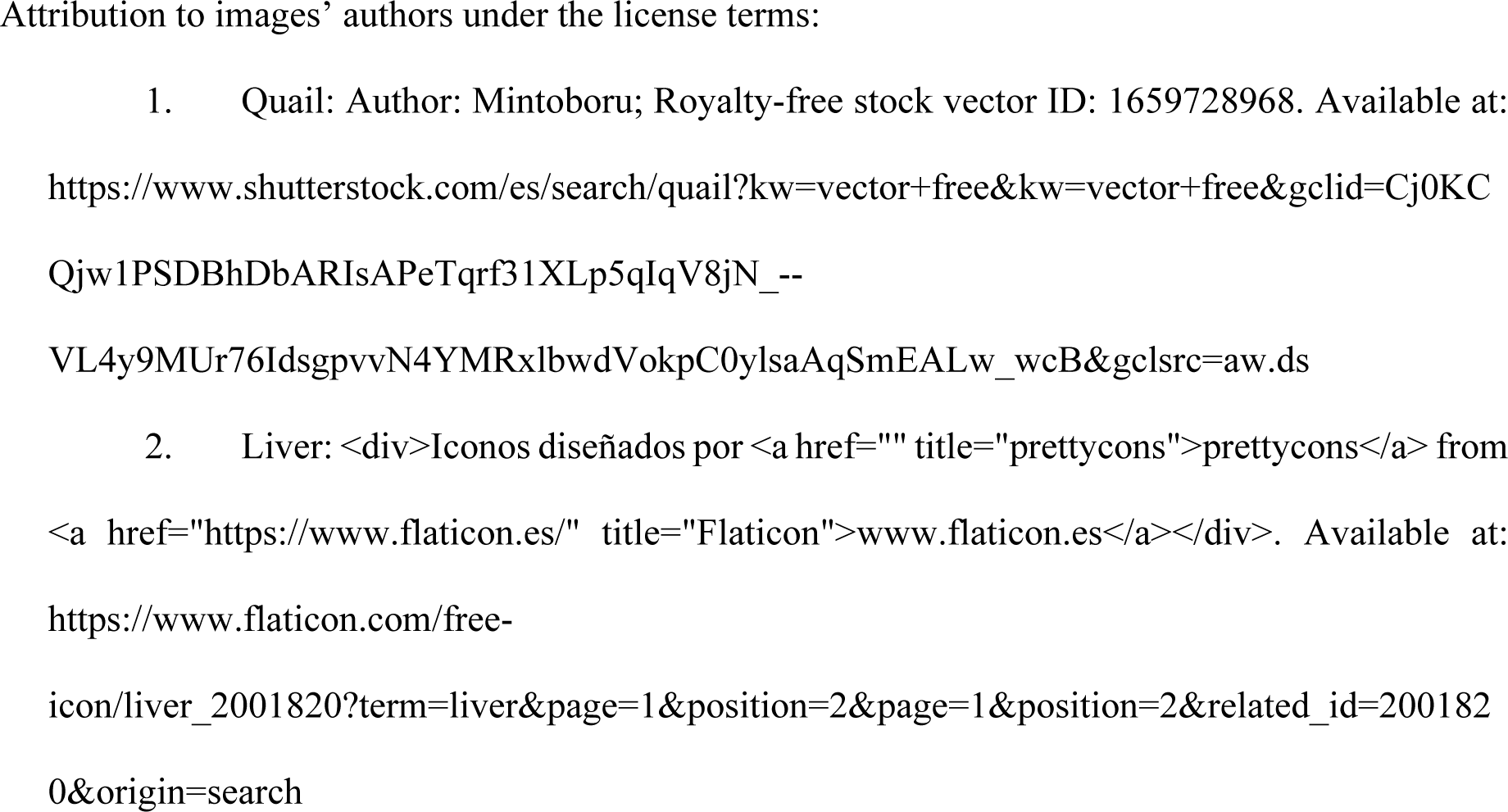
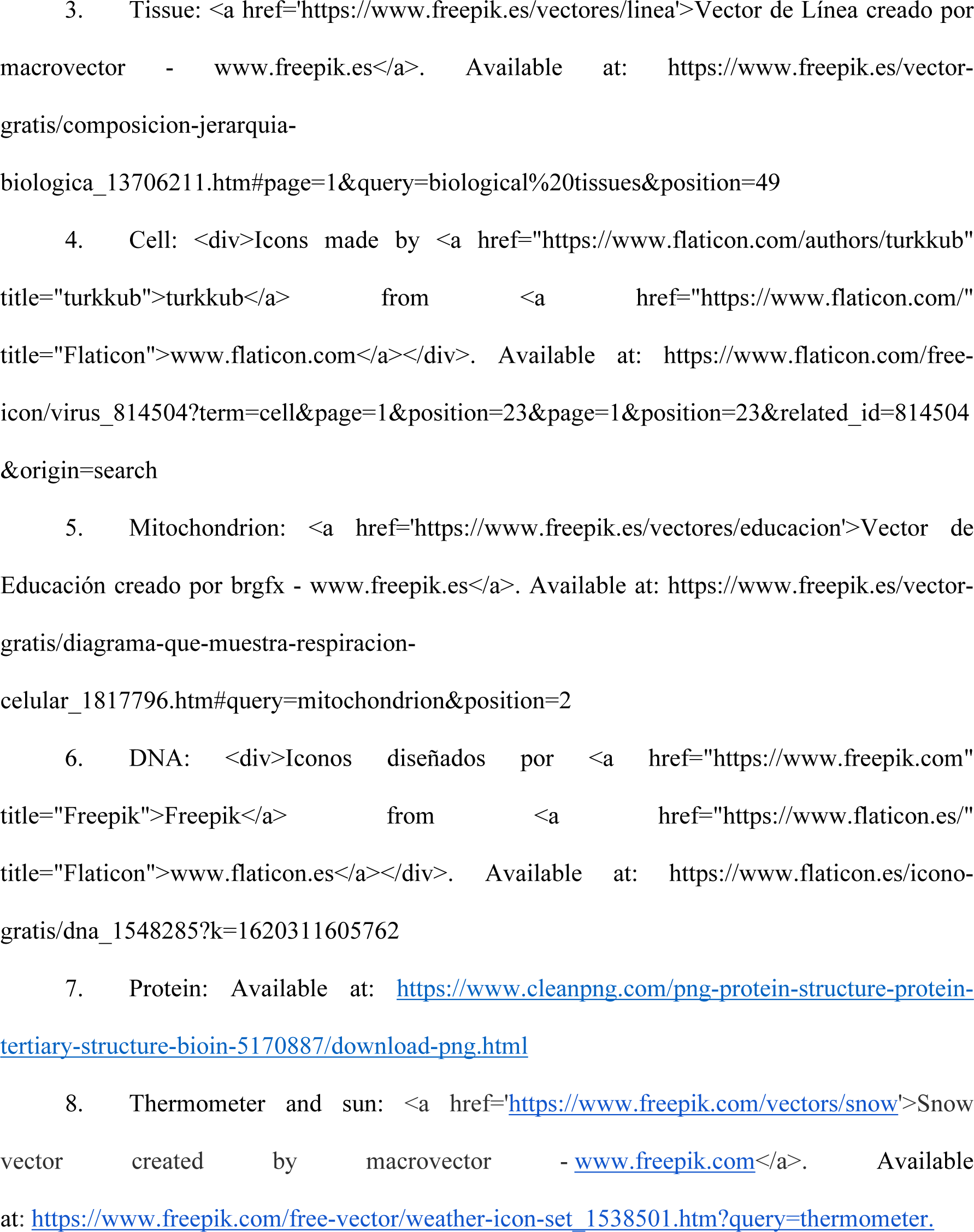

## Supporting information

S1 Appendix. Dietary supplementation. S2 Appendix. Heat stress protocol.

S3 Appendix. Assessment of female somatic maintenance traits. S4 Appendix. Assessment of egg production traits.

S5 Appendix. Assessment of oxidative stress and antioxidant system response.

S6 Appendix. Univariate statistical analysis. Contains Supplementary Tables S1-S4 and Figures S1-S4.

S7 Appendix. Supporting analysis of multivariate statistics.

